# Structural and chemical properties of insect’s chitin-containing extracellular matrices

**DOI:** 10.64898/2026.04.18.718486

**Authors:** Mario Wegmann, Marius Beck, Stefan Cord-Landwehr, Bruno Moerschbacher, Ernesto Scopolla, Carolin Fischer, Luca Bertinetti, Yael Politi, Hans Merzendorfer

## Abstract

Insect’s body barriers rely on specialized extracellular matrices that protect against harmful environmental influences. The outer barrier is the cuticle, which is composed of chitin, cuticle proteins and lipids. The peritrophic matrix (PM) serves as an inner barrier lining the midgut epithelium. It is composed of chitin fibers that are organized by PM proteins. While cuticle and PM proteins have received considerable attention in the past, supramolecular organization and physicochemical properties of the chitin component - particularly of the PM - remain poorly understood. Here, we combine synchrotron-based X-ray diffraction data from the PMs of lepidopteran and coleopteran insects with RNA interference (RNAi), mass spectrometric and histochemical analyses of the PM from *Tribolium castaneum* to determine chitin’s allomorphic state and degree of acetylation. The chitin of the PM exhibits signatures characteristic of dihydrate β-chitin along the entire midgut. In contrast, the cuticle is made of tightly packed α-chitin nanofibrils. Mass spectrometry revealed that the PM’s chitin is highly acetylated (>95%). RNAi silencing of gut-specific genes encoding chitin deacetylasesTcCDA6-9 further increases the degree of acetylation. Histochemical analyses staining chitin with different degrees of acetylation confirm the predominance of highly acetylated chitin in the PM. Notably, the larval cuticle has a layered organization with deacetylated chitin present in exo- and highly acetylated chitin in endocuticles. Depletion of both TcCDA1 or TcCDA2 impairs chitin deacetylation, which indicates that both proteins cooperate in their activity in the integument. These results establish fundamental principles of polysaccharide-based extracellular matrices, with broad implications for insect biology.

## Introduction

Chitin-containing extracellular matrices are fundamental structural components in insects, forming the basis of the cuticular exoskeleton, peritrophic matrix (PM) in the gut, and other protective barriers. These matrices not only protect insects from physical damage and desiccation but also serve as barriers against pathogens. While the structure and function of the cuticle have been extensively studied over the past decades [1], knowledge about the PM remains comparatively limited. The PM functions as a size-selective barrier that ensures efficient nutrient transport while simultaneously protecting the epithelium from mechanical damage, pathogens, and toxins [2-4]. It is composed of chitin fibrils and chitin-binding PM proteins (PMPs) that organize the PM into a 3D meshwork of chitin fibers and (glyco)proteins. Despite extensive characterization of some PM proteins, the molecular organization and physicochemical properties of the PMs chitin constituent remain insufficiently understood, particularly with respect to its crystalline allomorph and supramolecular packing. In the red flour beetle, *Tribolium castaneum*, it was shown that two chitin binding PMPs are required to maintain the structural integrity and size-selective permeability of the PM. RNAi-mediated silencing of the genes encoding these PMPs disrupts the midgut barrier, leading to abnormally high permeability, developmental arrest, and lethality [5]. How the PMPs organize the chitin fibers is largely unexplored. However, the physicochemical properties of chitin and its interaction with PMPs are thought to be influenced by the degree of polymerization (DP), the degree of acetylation (DA) and possibly by the patterns of acetylation (PA) [6-8].

Although the general composition of the PM is well established, major gaps remain in our understanding of its structural organization at the molecular level. Chitin is a linear polymer of *N*-acetylglucosamine (GlcNAc), which is produced chitin synthase (CHS). Insect genomes harbor two *CHS* genes, which are expressed in ectodermal epithelia like the epidermis (*CHS1*) to produce cuticle or endodermal epithelia like the midgut epithelium to produce PM (*CHS2*). CHS2 is present at the apical tips of midgut epithelial microvilli [9]. Chitin can be cleaved by various types of chitinases (CHT) or deacetylated to various degrees by chitin deacetylases (CDA) to eventually yield chitosan, a linear polymer of glucosamine (GlcN). The finding of CHTs and CDAs that are specifically expressed in the gut suggests that these enzymes may control some properties of the PM [10,11]. Chitin is thought to occur in three crystalline allomorphs – α-, β-, and γ-chitin – which differ in the orientation of their polymer chains and the pattern of hydrogen bonding between them [12]. α-Chitin, characterized by tightly packed antiparallel sugar-chains and dense intra- and intermolecular hydrogen bonding, is the predominant form in the cuticular exoskeleton of insects and other arthropods. It provides high mechanical strength and structural rigidity but also flexible and elastic properties, depending on the type of proteins that organize the cuticular matrix [13]. β-chitin, defined by parallel chains, high binding capacity for molecules, and less compact packing, exhibits enhanced flexibility and hydration, and is commonly observed in skeletal elements of marine organisms such as squid pens or the sheaths of tubeworms [14]. γ-Chitin is considered to be a mixed form with uneven number of parallel and antiparallel chains. It was initially suggested as a component of some peritrophic matrices [15], and has since been reported in the cuticle of silk glands and spinning ducts, as well as in specific insect cocoons or silk matrices [16]. Some midgut secretions of figwort weevils *Cionus scrophulariae* and *Cleopus pulchellus* have been identified as β-chitin [12,15]. Nevertheless, to date the PM’s chitin allomorph is poorly characterized in most insect species and may differ between different groups. Given the functional requirements of the PM as a flexible, hygroscopic, permeable, and continuously renewed structure, we hypothesized that PM chitin differs from cuticular chitin not only in its biochemical modifications but also in its crystalline and fibrillar organization.

Many insects express different sets of CHTs and CDAs in chitin-secreting tissues [1], which suggests that chitin occurs with different DP and DA depending on the tissue type. Based on the different localization of two CDAs in the locust cuticle, it was speculated about the possibility of different degrees of acetylation in the cuticles from fore- and hindgut [17]. The changes of the polymer’s properties and binding specificities associated with chitin deacetylation are significant as it increases the polymer’s solubility and flexibility and gives it a positive charge at neutral pH [18,19]. Hence, it is likely that the binding of cuticular proteins and PMPs is also affected by chitin deacetylation.

To date, it is unclear which chitin allomorph is found in the PM in insects, to which extent the PM’s chitin is deacetylated and whether the allomorphic state and the DA vary across different gut regions or between insect species. One of the major limitations is the lack of high-resolution structural data that can discriminate between different chitin allomorphs *in situ*. Mass spectrometry approaches (e.g. MS/MS of chito-oligosaccharides) have provided valuable composition-level information on patterns and the DA in chitinous material [20,21]. However, chitin allomorphs are defined by polymer chain packing and crystalline lattice parameters, which are determinable mainly by X-ray diffraction or solid-state NMR techniques (XRD, ssNMR) rather than by mass spectrometry [22,23]. Whereas these physical methods have been applied to analyze β-chitin from tubeworms, squid pens and cuttlefish bones [24-26], they have not been employed to study insects’ midgut crystallography since pioneering work in late 1970-1980 by Rudall and colleagues.

In this study synchrotron-based XRD experiments were performed on larval PM samples from *M. sexta* and *Z. morio* to compare lepidopteran and beetle species. We used mild extraction and purification methods, to minimize sample preparation artefacts. These analyses revealed that the PM of both species is predominantly composed of dihydrate β-chitin irrespective of the region within the midgut from which they originate. In *T. castaneum*, a tenebrionid beetle closely related to *Z. morio*, we could further show that the total mass fraction of chitin in the larval gut is very low and the DA is very high. The total mass fraction of polymerized GlcN and GlcNAc is less than 1% (w/w) and the DA slightly more than 95%. This high DA further increases to almost 98% after RNAi-mediated silencing of the genes encoding gut specific group IV chitin deacetylases *TcCDA6-9* in larvae of *T. castaneum*. Notably, the exocuticle of the larval body wall appears to have a low DA while the endocuticle has a high DA. Collectively, these findings highlight the need for an integrative analysis of chitinous matrices in multiple species to improve our understanding on the structural organization of chitinous biocomposites in general and of the insect PM in particular.

## Materials and Methods

### Chemicals

All standard chemicals were obtained in p.A. quality from Roth (Karlsruhe, Germany) and Serva (Heidelberg, Germany).

### Insect rearing

*Manduca sexta* larvae were obtained from a culture raised in the lab of Anke Steppuhn (Molecular Botany, University of Hohenheim, Stuttgart), kept at 27°C and 50% humidity and fed on standard artificial diet [27]. *Zophobas morio* larvae were bought at a local pet shop and kept on oatmeal until preparation.

The GA-1 strain of *T. castaneum* was used in all experiments [28], and cultured in the lab of HM since 2009. The beetles were kept on whole-wheat flour (Kaufland Bio, Neckarsulm, Germany) under standard conditions at 30°C and 50% humidity as described previously [29]. For experiments, larvae with a mean weight of 1.5 mg ± 0.1 mg were selected for dsRNA injection and grown on whole wheat flour, and the larvae were maintained under standard conditions for six days.

### Sample preparation for XRD

All insect larvae were anaesthetized on ice and decapitated prior to dissection. For regional PM analyses, anterior, median, and posterior midgut sections were dissected at first from ten *M. sexta* fifth instar larvae, and then the PM was isolated separately from each region. For whole-PM analyses, PMs were extracted either from six *M. sexta* fifth instar larvae or from twenty *Z. morio* sixth instar larvae. We focused on these species because their size allowed us to extract the PM from the midgut in amounts required for XRD measurement.

Isolated PM samples were repeatedly washed in phosphate-buffered saline (PBS) until visually transparent to remove residual gut contents. Subsequently, samples were washed with deionized water to remove salts. PM samples were either lyophilized for standard XRD analysis. For XRD measurements, PM samples were mounted on silicon nitride frames and overlaid and sealed with 12.5 µm Kapton® foil, to ensure stable mounting.

### XRD measurements

XRD measurements were performed at the µSpot beamline of BESSY II (Berlin, Germany) in transmission geometry. The incident X-ray beam energy was set to an energy of 18 keV (λ = 0.689 Å) using a multilayer monochromator. At the sample position, the beam was focused to a spot size of approximately 100 µm, and a 250 µm beamstop was used to block the direct beam. Diffraction patterns were collected at a sample-to-detector distance of 340 mm, calibrated using quartz powder. This configuration provided access to a scattering vector range (q-range) of around 0.1–40 nm^−1^, covering most reflections characteristic of known chitin allomorphs. Data were recorded using a two-dimensional Dectris Eiger 9M detector (2070 × 2167 pixels). To minimize radiation damage, preliminary beam-damage tests were performed on sacrificial regions of the samples by varying acquisition time and step size. Based on these tests, all mapping scans were conducted with a step size exceeding the beam footprint, ensuring that each sample region was exposed only once. The acquisition time was set to 60 s per measurement position.

### XRD data processing and analysis

Collected two-dimensional diffraction patterns were corrected for detector geometry and background scattering and subsequently azimuthally integrated to generate one-dimensional intensity profiles as a function of the scattering vector q. Peak positions were extracted from the resulting profiles, fitted and averaged over the scanned regions using custom analysis scripts. Representative fits are shown in Figures S3 and S4.

### Quantitative enzymatic-mass spectrometric (QEMS) analysis of chitinous polymers

For quantitative analysis of chitinous polymers, gut tissues were dissected from *T. castaneum* larvae, as these beetles are amenable to systemic RNA interference (RNAi) experiments. For this purpose, 1.5 mg larvae were injected with dsRNA containing solutions to silence *TcVER, TcCDA6, TcCDA7, TcCDA8*, or *TcCDA9* genes as described below. Each experimental group comprised ten pooled guts. The dissected tissues were immediately frozen, lyophilized, and subsequently shipped to the University of Münster for quantitative enzymatic–mass spectrometric (QEMS) analysis as described previously [30]. In brief, the chitinous polymer fraction was isolated from the lyophilized material and subjected to isotopic *N*-acetylation to label deacetylated GlcN residues. The isotopically modified polymers were enzymatically hydrolyzed to monomeric constituents, which were quantified by mass spectrometry using a double-isotopically labeled internal standard ([^13^C_2_, ^2^H_3_]-*N*-acetyl-D-glucosamine).

Quantification of the native and isotopically labeled monomers was performed via LC-MS using a Synapt XS HDMS 4k mass spectrometer (Waters; Milford, MA, USA) to have high resolution and high sensitivity to enable the determination of the degree of acetylation and the total content of chitinous polymers in each sample. The separation of monomers was performed using an Acquity Premier UPLC System (Waters, Milford, MA, USA) using an Acquity UPLC BEH Amide column (1.7 μm, 2.1 mm × 50 mm; Waters, USA) with a VanGuard precolumn (1.7 μm, 2.1 mm × 5 mm; Waters, Milford, MA, USA) and a column oven temperature of 40°C. The flow rate was set to 0.4 mL/min and the separation was performed over 8.0 min with the following gradient elution profile: 100% (v/v) A (80:20 ACN:H_2_O with 10 mM NH_4_HCO_2_ and 0.1% (v/v) formic acid) for 0.0-3.0 min, linear gradient to 80% (v/v) B (20:80 ACN:H_2_O with 10 mM NH_4_HCO_2_ and 0.1% (v/v) formic acid) from 3.0-6.0 min, constant 80% (v/v) B from 6.0-6.5 min, linear gradient to 100% (v/v) A from 6.5-6.7 min and then 100% (v/v) A for 6.7-8.0 min to equilibrate the column for the next run. For the detection of the chitin monomers using the Synapt XS HDMS 4k mass spectrometer (Waters, Milford, MA, USA) following parameters were used. We used positive sensitivity mode with a mass range of m/z 50 to 1200 and a scan time of 1.0 s. As internal calibration solution leucine encephalin (100 pg/ µl) was injected in intervals of 10 s for 1 s as lock spray calibration. The source capillary voltage was set to 0.5 kV, sampling cone to 10, and source offset to 4. The source was set to 60°C and the desolvation temperatures to 150°C. Flow rates of 0 L/h cone gas and 500 L/h desolvation gas were used, and the nebulizer pressure was set to 6.5 bar.

### dsRNA synthesis and injection

cDNA fragments for *TcVER* (encoding the *vermillion* gene, positive control for RNAi), *TcCDA1* (EU019711.1), *TcCDA2* (EU019713.1), *TcCDA6* (EU190489.1), *TcCDA7* (EU190490.1), *TcCDA8* (EU190491.1) and *TcCDA9* (EU190492.1) were amplified using gene specific primers (Table S1). The PCR products were ligated into the pGEM-T-easy vector (Promega, Mannheim, Germany), and the resulting plasmids were used as templates for a PCR with sequence-specific primers containing a T7 promoter sequence at their 5′ ends. The PCR products were subjected to agarose gel electrophoresis, excised and purified with help of the QIAquick Gel Extraction Kit from Qiagen (Hilden, Germany). The PCR fragments were then used as a template for dsRNA synthesis, which was performed with the HiScribe^®^ T7 High Yield RNA Synthesis Kit (New England Biolabs). After hybridization of the complementary ssRNAs, the resulting dsRNA was purified via phenol-chloroform-isoamylalcohol extraction and precipitated with ammonium acetate. Injection of 400 ng dsRNA into 1.5 mg mid-sized fourth instar larvae was performed as reported previously [31]. Total RNA was prepared and transcribed into cDNA (see below). RNAi efficiency was determined by qPCR and only statistically significant knockdowns below 80% were further analyzed.

### qPCR analysis

Total RNA was prepared from pools of 3 whole larvae of *T. castaneum* four days after dsRNA injection using the RNeasy Mini Kit (Qiagen, Hilden, Germany) followed by DNaseI digestion (RNase-free, Thermo Scientific, Waltham, USA). cDNA synthesis was performed using the biotechrabbit cDNA Synthesis Kit and oligo(dT) primers following the manufacturers’ instructions (biotechrabbit GmbH, Berlin, Germany). The qPCRs were performed in triplicates with the Magentic Induction Cycler (Fujirebio) using qPCR SyGreen Mix Fluorescein (Nippon Genetics, Tokyo, Japan), 100 ng of the respective cDNA and pairs of gene specific primers (Table S1). Relative gene expression was calculated based on the comparison of C_T_ values for the gene of interest and the reference gene *TcRPS6*. The specificity of the PCR was confirmed by melting-curve analysis and mean normalized expression was determined according to Simon [32]. RNAi efficiencies for silencing the genes *TcCDA1-2* and *TcCDA6-9* are given in Fig. S1.

### Cryosectioning

Cryosections of larvae were performed using a modified protocol [33]. The larvae were anesthetized on ice for 5 min and then decapitated. The larvae were incubated in 4% (w/v) paraformaldehyde rotating for one hour at room temperature. The samples were washed 10 min in 1x PBS and then dehydrated by a sucrose gradient from 10% (2x 10 min at RT and rotating), 20% (15 min at RT) and 30% (1h at RT). After that the larvae were embedded in Tissue-Tec (Leica, Wetzlar, Germany) and frozen at -20°C. Transversal cryosections were generated as described previously using a Leica cryostat (CM1850, 160 mm steel blade with C profile, 15 µM slice thickness) and SuperFrost Plus microscope slides to collect the sections [5]. The resulting specimens were dried at 40°C on a hotplate for tissue attachment. Finally, the samples were stained with fluorescent dyes, covered by VectaShield mounting medium (Vector Labs Inc., Burlingame, USA), the open sides were sealed with nail polish, and the sections were viewed under a fluorescence microscope (Zeiss Axio Observer V7, Oberkochen, Germany) using appropriate filter sets.

### Histochemical analysis

For histochemical characterization, the tissue sections were subjected to fluorescence staining to visualize specific structural components. Cell nuclei were stained with DAPI (4′,6-diamidino-2-phenylindole). To label high DA chitinous structures, a chitin**-**binding domain from a *Bacillus licheniformis* chitinase coupled to superfolder GFP (CBD-sfGFP) was used [34]. In addition, a chitosan affinity protein from an enzymatically inactivated *Bacillus sp*. chitosanase fused to the red fluorescent protein mKATE (CAP-mKATE), was used to specifically label low DA chitin [35]. Both fusion proteins were obtained from the lab of Bruno Moerschbacher, University of Münster.

## Results and Discussion

### X-Ray Diffraction of whole PM samples indicates the presence of beta chitin

To assess the structural organization of chitin in isolated PM, XRD analyses were performed on freeze-dried PM material extracted from the entire midgut of *M. sexta*. To avoid preparation artefacts leading to β → α recrystallization of chitin, no protein extraction was performed on the isolated PM. The diffraction profiles of whole-PM preparations closely matched those obtained from individual midgut regions, confirming that regional analyses reflect the global structural organization of the PM. Whole-PM diffraction profiles exhibited broad and poorly resolved reflections consistent with semi-crystalline chitin and the presence of proteins. We identified the di-hydrate form of beta-chitin, implying that freeze-drying did not remove bound water, or that these were re-intercalated during further sample preparation.

As a reference for α-chitin in its native state, the XRD profile of the *T. castaneum* elytral cuticle was included. This α-chitin profile exhibits sharper and more intense Bragg reflections with well-resolved (020), (110), and (013) peaks, reflecting higher crystallinity. In particular, the α-chitin sample displays a prominent low-q correlation peak at q ≈ 1.36 nm^−1^ (corresponding to ∼6.3 nm d-spacing), which is a well-known feature of tightly packed chitin nanofibrils ([36], Fig. 2A, Tab. 1). This feature is practically absent in PM samples of both *M. sexta* and *Z. morio*, which instead show only a broad, weak maximum in this region. This suggests a looser and disordered nanoscale arrangement of the fibrils, consistent with the more hydrated, gel-like PM matrix rather than a tightly packed fiber composite. Furthermore, peak shapes in the PM are broader and less distinct than in the α-chitin control, indicating smaller coherent domain sizes and increased structural heterogeneity.

**Table 1:** Empirical peak separation values for three crystalline chitin allomorphs (α- and β-chitin) reported in literature. α-Chitin exhibits a smaller separation between the (110) and (020) peaks (∼7.06–7.12 nm^−1^), whereas β-chitin shows a larger separation between the (110) and (010) peaks (∼7.79–8.04 nm^−1^). For our PM samples, Δq ≈ 8.1–8.3 nm^−1^ consistently exceeds the typical α-chitin range and instead falls in the β-chitin range, supporting the conclusion that PM chitin has a β-chitin–like lattice structure.

| α-Chitin$\Delta q$ [(110)–(020)] ($nm^{-1}$) | β-Chitin$\Delta q$ [(110)–(010)] ($nm^{-1}$) | Reference |
| --- | --- | --- |
| 7.08 | 8.04 | [45] |
| 7.12 | 7.79 | [16] |
| 7.06 | 7.90 | [44] |

### XRD analyses reveals no major differences of chitin allomorphs in the PM from different midgut regions

To investigate the structural organization of chitin within the PM, XRD measurements were performed on PM samples isolated from the anterior, median, and posterior midgut regions of fifth instar *M. sexta* larvae. All diffraction profiles exhibited clear but broad chitin-associated reflections, indicative of low crystallinity. Across all regions, signals were consistently detected at q-values of 5.7 (± 0.07) nm^−1^, 14.10 (± 0.20) nm^−1^, and 19.00 (± 0.15) nm^−1^, corresponding to the characteristic β-chitin reflections (010), the combined (100)/(020) and (110), and the group (013)/(112)/(121), respectively in its di-hydrated form (Fig. 1). No systematic differences in peak position or width were observed between anterior, median, and posterior PM samples (Fig. S3). This is further supported by Gaussian peak fitting of the diffraction profiles, which confirms the consistency of peak positions and relative contributions across regions and replicates (Table S2). Occasionally occurring sharp spikes detected in individual measurements (e. g. at q ≈ 16.5 nm^−1^ in anterior PM sample B) are due to contamination of the sample holder or buffer salts and were absent in corresponding reference measurements (Fig. S2), confirming that these signals are unrelated to the PM’s chitin structure.

**Figure 1.**
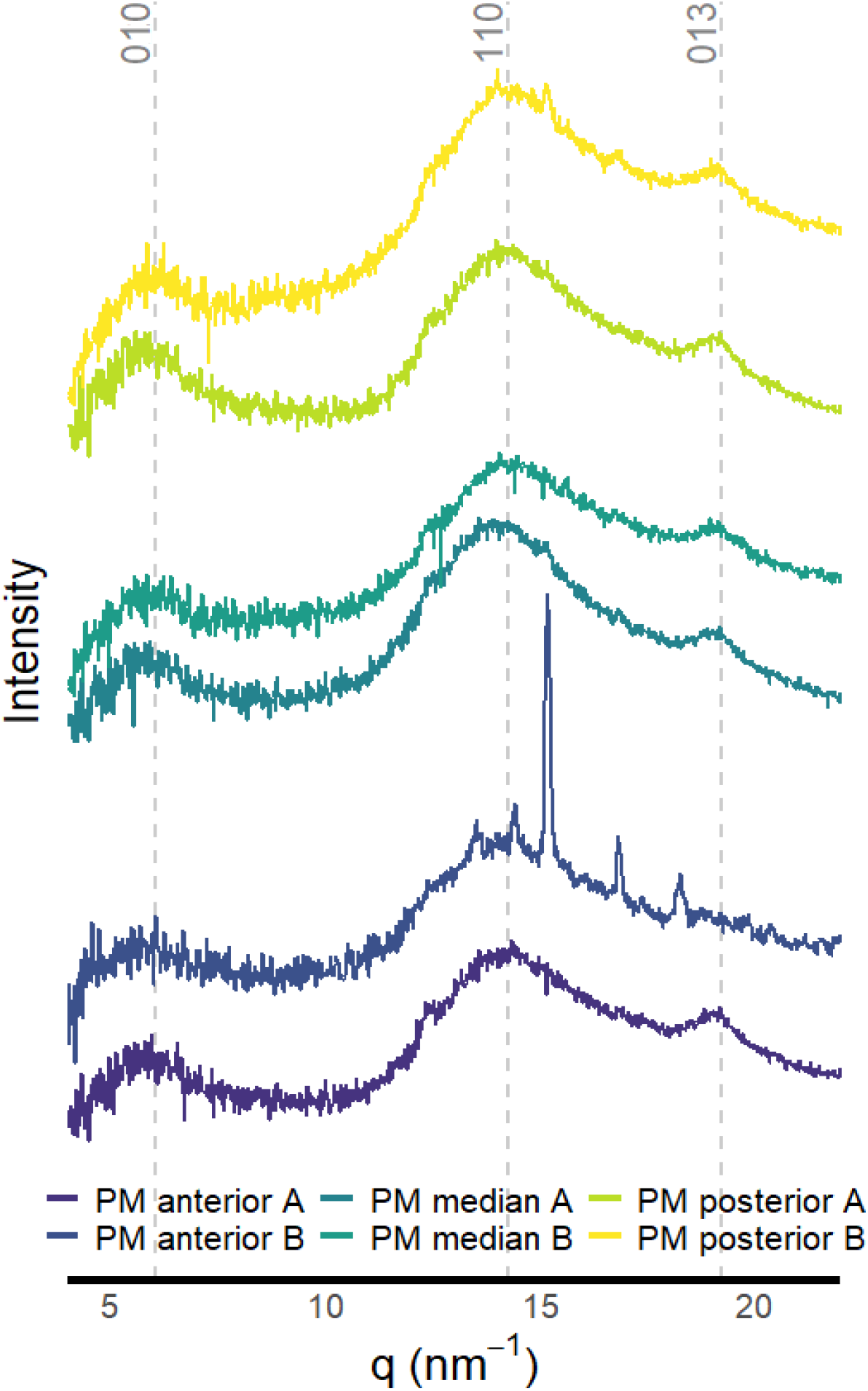
Azimuthally integrated X-ray diffraction profiles of freeze-dried peritrophic matrix (PM) from different midgut regions of *Manduca sexta*. Representative, azimuthally integrated 1D diffraction profiles of a freeze-dried PM samples from different midgut regions of *Manduca sexta*. Shown are two replicates per region, from the anterior, median and posterior midgut. The intensity (arbitrary units [A. U.]) versus q (nm-1) profiles show a single peak in the low-q range (020) and several peaks in the high-q range (110) and (013).

The high degree of overlap between diffraction profiles from different midgut regions demonstrates that chitin organization within the PM is structurally uniform along the anterior– posterior axis. Consequently, regional differences in PM properties such as permeability or exclusion size, as they were observed in *T. castaneum* [5], are unlikely to arise from changes in chitin allomorph and more plausibly reflect differences in PM thickness, chitin–protein crosslinking, or interactions with regionally expressed PM proteins.

### XRD profiles indicate high similarities of the PM’s chitin structure in different species

To determine whether the β-chitin allomorph of the PM is conserved beyond lepidopteran species, whole-PM samples from *M. sexta* were compared with those from the coleopteran *Z. morio*. The darkling beetle was selected to allow comparison with the established beetle model system *T. castaneum*, which served as the basis for complementary biochemical and histochemical analyses in this study. The diffraction profiles obtained for the *Z. morio* PM closely resembles those obtained from *M. sexta*, displaying comparable peak positions for all chitin-associated reflections (Fig. 2). Gaussian peak fitting further confirms the similarity in peak positions and structural contributions between species (Fig. S4, Table S3). While an increased relative intensity of the (110) reflection compared to the (010) reflection was observed in *Z. morio*, no shifts in peak position were detected. This difference could reflect species-specific variations in crystallite size, orientation, or protein–chitin interactions.

**Figure 2:**
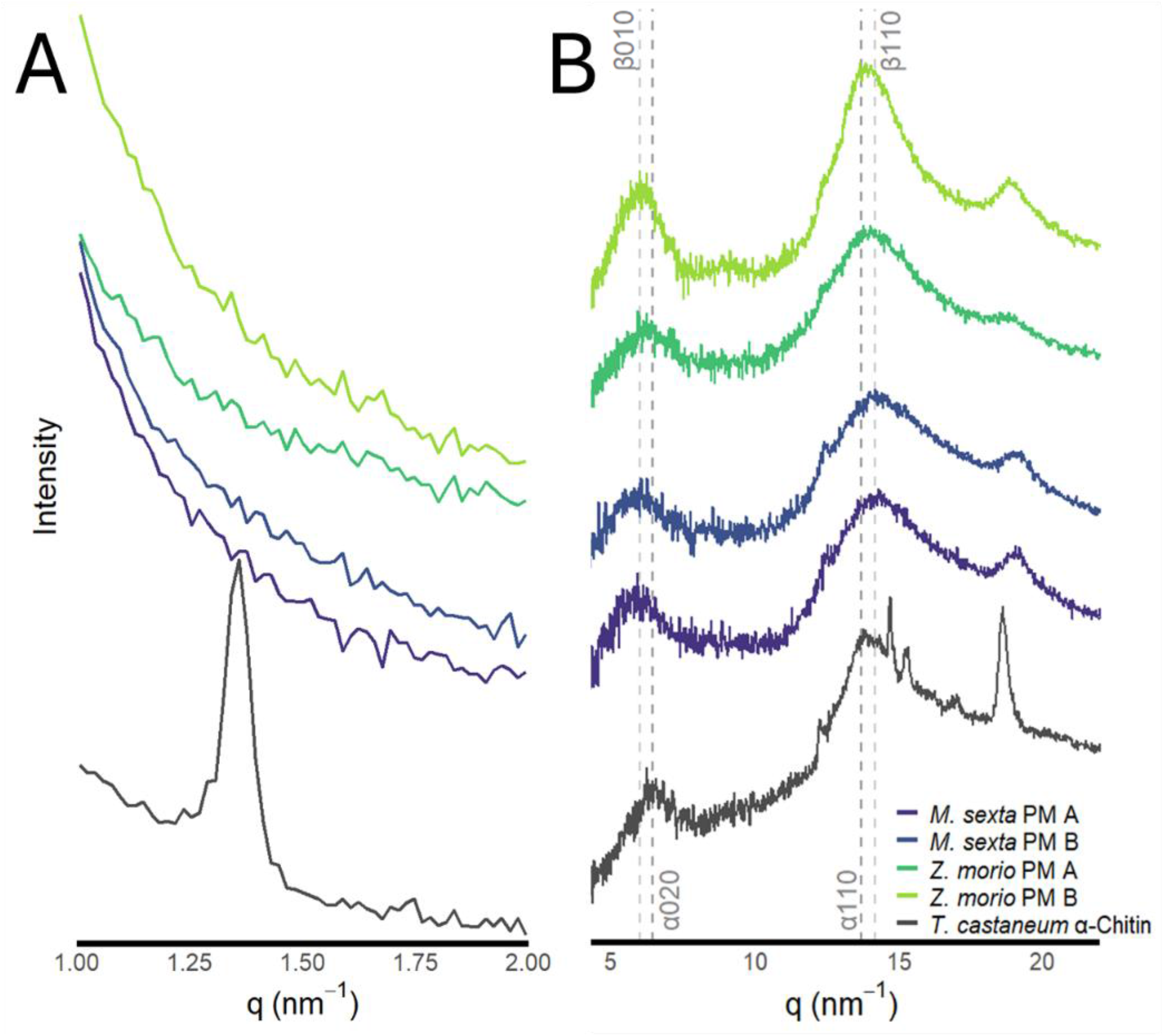
Structural comparison of PM chitin with highly ordered α-chitin. Representative, azimuthally integrated 1D diffraction profiles of freeze-dried PM samples from whole midgut PM of *M. sexta* and *Z. morio*. Shown are two replicates per species. The profile obtained for the pupal elytra from *T. castaneum* is shown as a α-chitin internal reference. **(A)** Low-q diffraction profiles emphasizing the first-order packing peak associated with chitin chain organization. All PM samples exhibit a broad low-q maximum, which corresponds to the fundamental packing distance of chitin fibrils and reflects limited long-range order and heterogeneous spacing within the PM. In contrast, the α-chitin of the cuticle reference displays a sharper and more intense packing peak, consistent with highly ordered crystalline packing of antiparallel chitin chains in α-chitin. **(B)** Wide-angle diffraction profiles comparing PM samples from *M. sexta* and *Z. morio* with highly crystalline α-chitin from *T. castaneum* elytra. The α-chitin reference exhibits sharp and well-resolved Bragg reflections corresponding to the (020), (110), and (013) lattice planes, indicative of a highly ordered orthorhombic α-chitin crystal structure. In contrast, the PM samples show broadened and less well-defined peaks, reflecting reduced crystallinity and increased structural heterogeneity. Notably, the increased separation between the (020) and (110) reflections in the PM samples suggests expanded interplanar spacing and a less dense lateral packing of chitin chains, likely arising from protein incorporation and functional adaptation of the PM.

Together, these results indicate that the chitin component of the PM in both *M. sexta* and *Z. morio* predominantly adopts a dihydrate β-chitin lattice organization. The conservation of this structural signature between a lepidopteran and a coleopteran species suggests that β-chitin is a common form in the PM of these large insect orders. It appears plausible that next to the presence of mucin-like PM proteins the enhanced hydration capacity associated with β-chitin adds a structural basis for the PM’s function as a hydrogel-like, permeable barrier within the digestive system, particularly in the anterior region [3].

### Midgut chitin is highly acetylated, and the DA is further increased in response to RNAi silencing midgut-specific chitin deacetylases

Following the structural characterization of the chitin allomorph in the PM, we next quantified the overall intestinal abundance and acetylation state of chitin. Quantitative enzymatic mass spectrometry (QEMS) was employed to determine the total amount of polymerized chitin in the midgut of *T. castaneum* and to determine the DA. The analysis revealed that chitin constitutes only a minor fraction of the midgut’s total mass. In control larvae injected with ds*TcVER*, polymerized GlcNAc accounted for about 0.57% (± 0.02%) of the midgut’s total mass, whereas GlcN represented only 0.02% (± 0.001%; Fig. 3A). These data demonstrate that the overall chitin content in the midgut, containing the PM, is low in comparison to other components like nucleic acids, proteins, glycoproteins and lipids [37]. Yet chitin has important functions as it acts as a structural scaffold within the PM.

**Figure 3.**
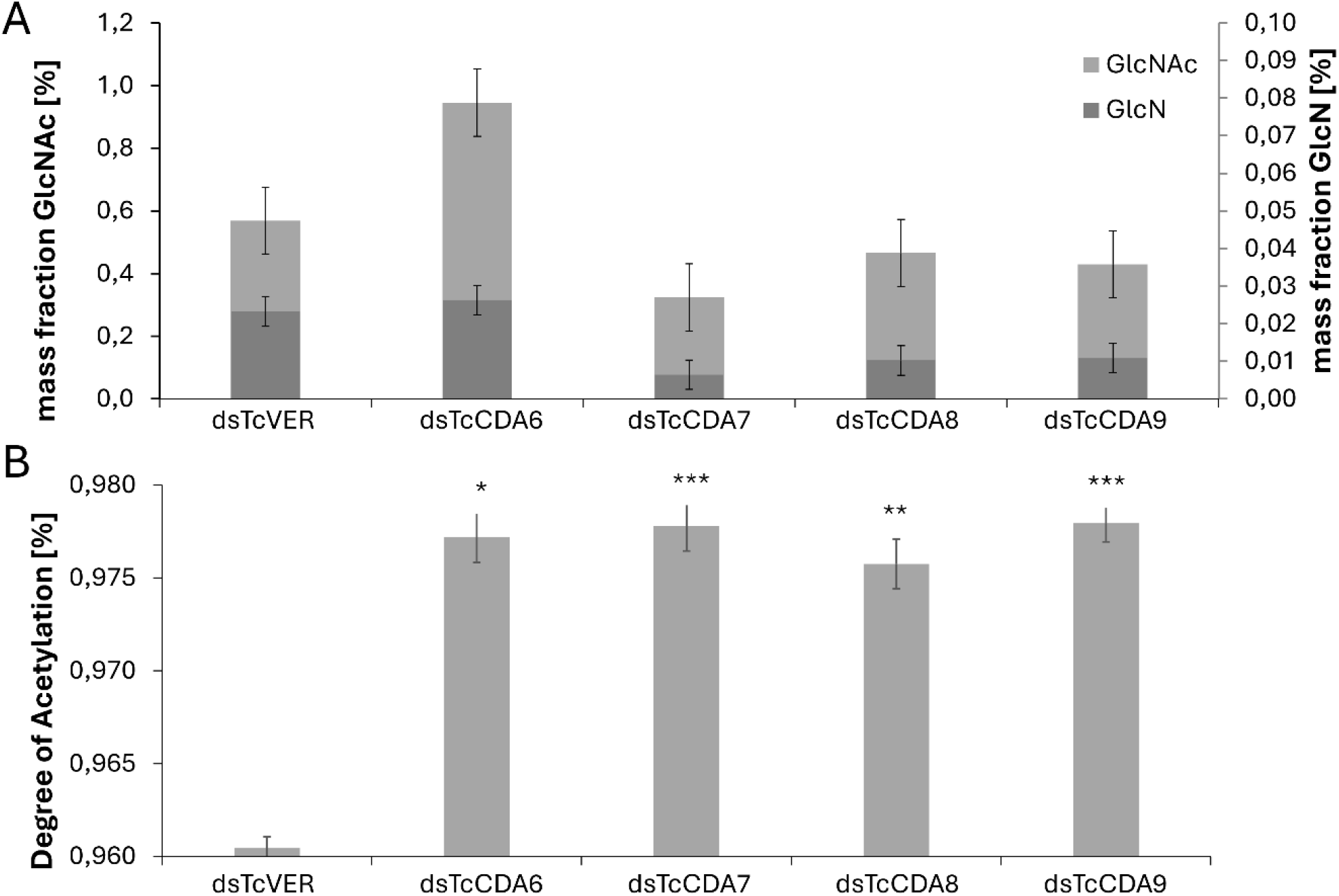
Low chitin abundance and high degree of acetylation in the midgut of *Tribolium castaneum*. Quantification of chitin-derived monosaccharides and degree of acetylation (DA) in midguts of larvae 4 days after dsRNA injection targeting *TcCDA6-9* genes. **(A)** Relative mass fractions (w/w) of GlcN [%] (dark grey) and GlcNAc [%] (grey) (± SD) in midgut samples for each RNAi treatment. **(B)** Degree of acetylation (DA [%]) in midgut samples following RNAi treatment. Bars represent mean ± SD of three replicates from one pooled sample (10 midguts per group). Statistical significance was assessed by unpaired two-tailed Student’s *t*-tests comparing each treatment group to the control (*dsTcVER*). Significance levels are indicated as: p < 0.05 (*),p < 0.01 (**),p < 0.001 (***).

In addition to measuring the midgut’s total chitin content, QEMS enabled the determination of the DA. In control larvae, the DA was 96.0% (± 0.3%) indicating that the vast majority of the midgut’s chitin is present in its acetylated form. However, there are four CDAs (*TcCDA6-9*) whose genes are expressed in the larval midgut of *T. castaneum* [38]. To assess the contribution of these CDAs to the deacetylation of the midgut chitin, RNAi-mediated knockdown of each of the *TcCDA6-9* genes was performed. In all cases, RNAi resulted in a further increase in DA, with values of 97.7%, 97.8%, 97.6% and 97.8% (± 0.1, 0.2, 0.2 and 0.2%) for *TcCDA6, TcCDA7, TcCDA8* and *TcCDA9*, respectively.

Statistical significance was assessed by unpaired two-tailed Student’s *t*-tests comparing each treatment group to the control (dsVER). Significance levels are indicated as: p < 0.05, ** p < 0.01, *** p < 0.001. (Fig. 3B).

Notably, the knockdown of none of these CDAs caused lethality or affected growth and development. These findings may indicate that midgut-specific CDAs are either non-essential, at least under laboratory conditions, or that they have redundant functions. However, the latter may be contradicted by the finding that simultaneous knockdown of *TcCDA6-9* in single larvae did also not reveal abnormal phenotypes (not shown). Together, our data show that midgut chitin is highly acetylated and that deacetylated chitin represents only a very minor fraction, even in the presence of multiple gut-expressed chitin deacetylases. Deacetylated chitin may be present in the PM but could also be associated with the cuticles of trachea supplying oxygen to the midgut epithelium [33].

### The PM consist of highly acetylated β-chitin while the α-chitin of the cuticle from the larval body wall exhibits layers of different DAs

To further assess the DA of chitin in the PM but also in the cuticle, we performed histochemical staining of transversal cryosections from larvae using two fluorescent probes with complementary binding specificities. Chitin with high DA was visualized using an sfGFP-tagged (green) chitin-binding domain (CBD) derived from a *B. licheniformis* chitinase, whereas chitin with low DA was probed using an mKATE-tagged (red) chitosan affinity protein (CAP) based on a catalytically inactivated *B. sp* chitosanase.

No consistent m-KATE-CAP signal (red) was detected in the midgut, the midgut lumen or within the peritrophic matrix, while the PM was clearly stained by the sfGFP-CBD probe (Fig. 4A). Therefore, we can conclude that the vast majority of the midgut’s high DA chitin is found in the PM. In line with the QEMS measurements of the DA of the midgut’s total chitin, we found no histochemical evidence for the presence of low DA chitin in the midgut, trachea or in the PM. Hence, deacetylated chitin represents only a minor fraction of total midgut chitin. Low-abundance or highly localized low DA chitin - e.g. restricted to discrete microdomains within the PM - may remain below the detection limit in cryosections and would require higher-resolution imaging or broader sampling across the midgut to resolve.

**Figure 4.**
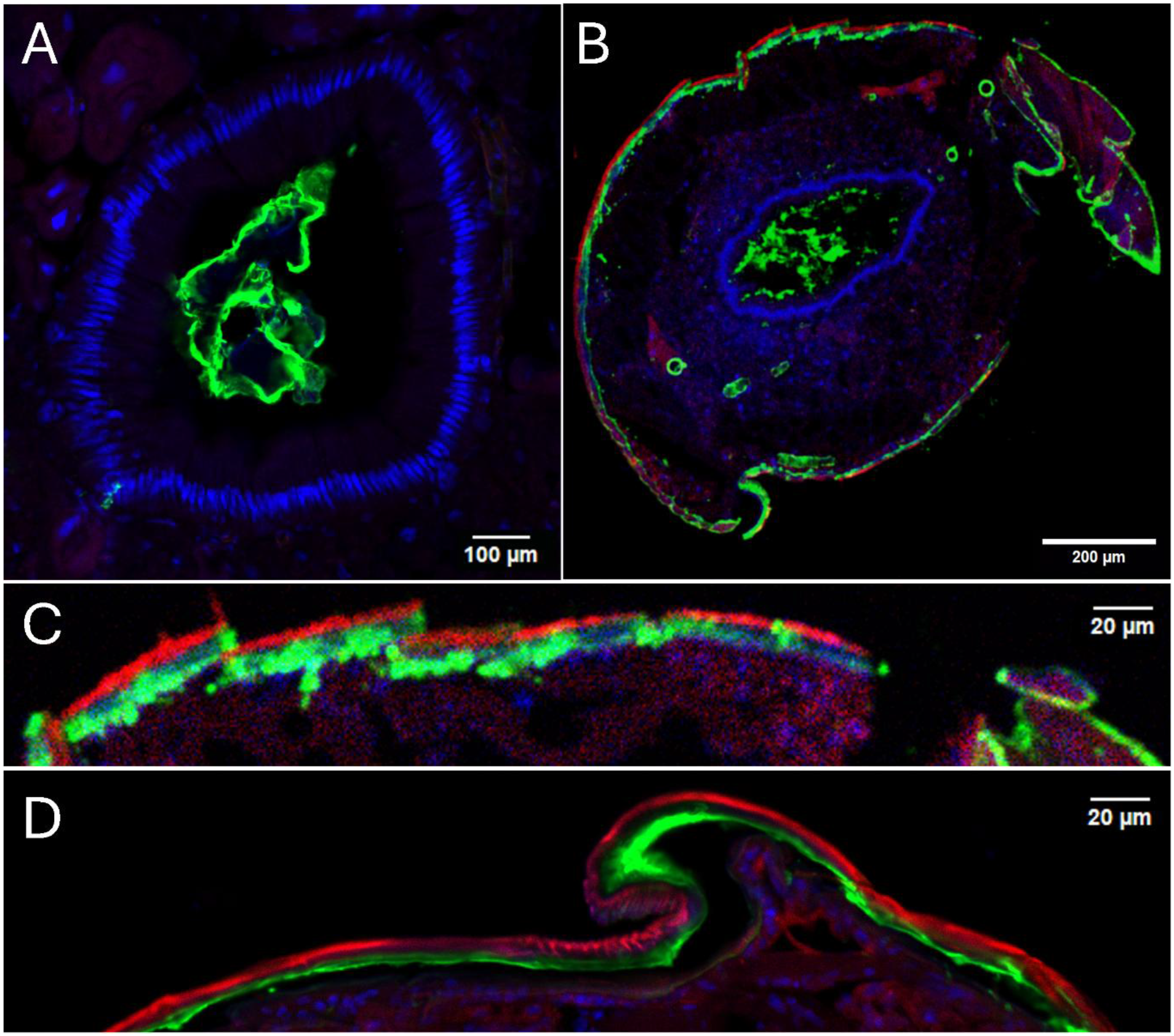
Spatial distribution of chitin and chitosan in the midgut and cuticle of *Tribolium castaneum*. Histochemical staining of cryosections derived from *Tribolium castaneum* larvae four days after *dsTcVER* injection. **(A)** Close-up of the midgut of a larva. Nuclei of midgut epithelial cells are stained with DAPI, and high DA chitin of the PM is stained with a CBD-sfGFP fusion protein. Notably, no staining is observed in the PM with CAP-mKATE protein indicating the absence of low DA chitin. **(B)** Transversal cryosection through a larvae depicting cuticle and the median midgut. High DA chitin is stained with CBD-sfGFP and low DA chitin CAP-mKATE protein. **(C)** Close-up on the apical cuticle taken from (B), where low DA chitin appears to present in the exocuticle and high DA chitin in the endocuticle. (D) Higher magnification image of another cuticle region from another cryosection stained again with CBD-sfGFP and CAP-mKATE protein.

In contrast to the midgut, a robust mKATE-CAP signal was consistently detected in the outer layers of the larval cuticle, whereas the sfGFP-CBD probe predominantly labeled inner cuticular layers (Fig. 4B–D). This staining pattern indicates a stratified organization of chitinous material in the larval cuticle, with preferential localization of low DA chitin in more external layers of the procuticle (exocuticle) and preferential high DA chitin in procuticular layers underneath (endocuticle). CDA activity has been shown to be essential for proper lamellar organization and higher-order assembly of the chitin matrix rather than for bulk chitin synthesis [37]. A specific distribution of chitin with different DA has been suggested so far only in *Locusta migratoria*, based on the presumed activities of two cuticle-associated CDAs acting on the cuticles of the fore- and the hindgut [17].

### Different DAs in the cuticle from the larval body of *T. castaneum* wall depend on the function of TcCDA1 and TcCDA2

To test the hypothesis that cuticle-associated chitin deacetylases contribute to the formation of the highly deacetylated outer cuticular layer, cryosections of larvae were analyzed following RNAi-mediated knockdown of *TcCDA1* and *TcCDA2* [11]. These CDA’s are known to be expressed in the epidermis and are essential for the higher-order organization of chitin fibers in the cuticle [38,39].

In *dsTcVER*-injected control larvae, a distinct mKATE-CAP-positive layer was consistently detected in the apical region of the cuticle, indicating the presence of chitin with a low DA (chitosan when fully deacetylated). Conversely, this signal was absent in both *TcCDA1-* or *TcCDA2-*injected larvae, while the sfGFP-CBD probe is still staining the underlying chitin layers with a high DA (Fig. 5).

**Figure 5.**
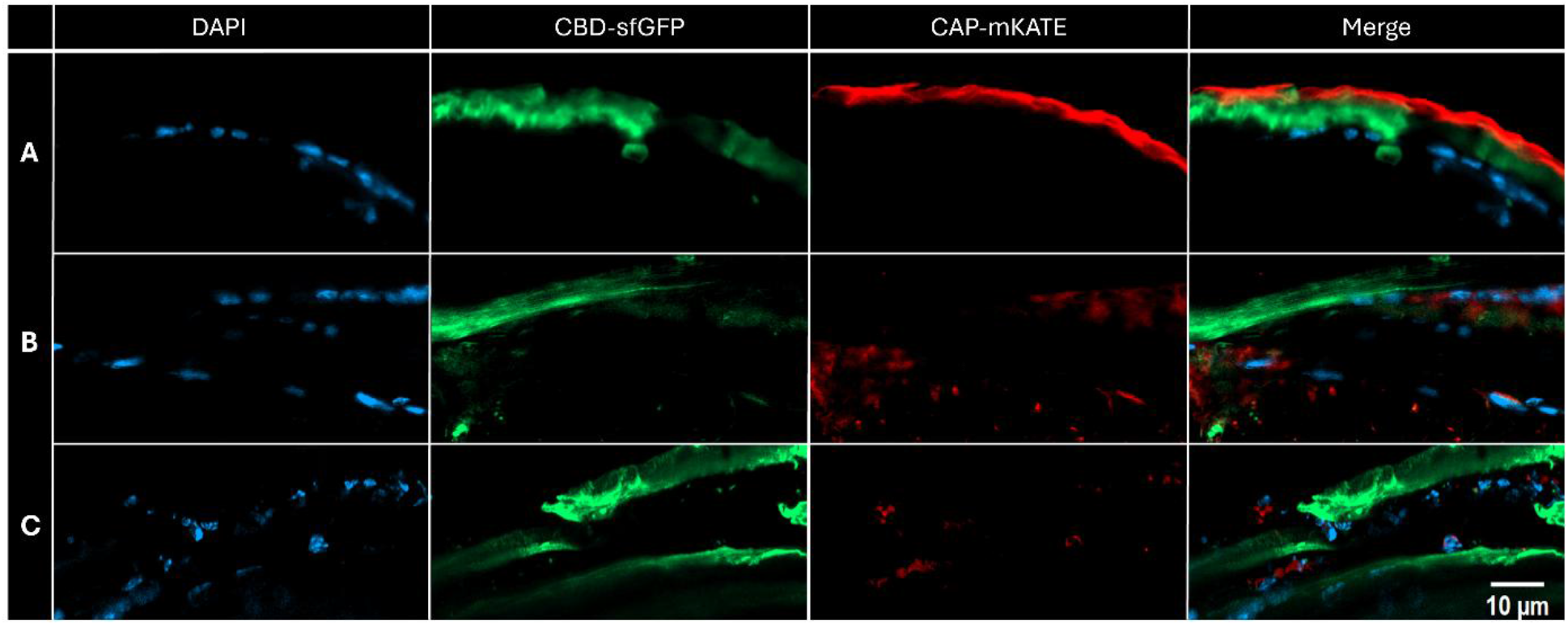
RNAi to silence TcDCA1 or TcCDA2 impairs chitin deacetylation in the exocuticle. Cryosections were derived from *T. castaneum* larvae 4 days after dsRNA injection. Nuclei of epidermal cells were stained with DAPI, high DA chitin is stained with a CBD-sfGFP fusion protein and low DA chitin with a CAP-mKATE fusion protein. **(A)** Close-up on the cuticle of a *dsVER*-injected control larva. High DA chitin is detected again in the endocuticle and low DA chitin in the exocuticle **(B, C)** Close-up on the cuticle of a *dsTcCDA1*-injected larvae (B) and a *dsTcCDA2* injected larvae (C). The staining of low DA chitin with the CAP-mKATE fusion protein in the exocuticle disappears in both cases.

Evidence from multiple insect systems indicates that Group-I chitin deacetylases are not fully redundant but may fulfill partially distinct functions, with CDA1 more closely associated with chitin deacetylation and CDA2 contributing to the structural organization of the cuticle [40,41, 17]. The finding that the knockdown of both TcCDA1 and TcCDA2 result in missing chitin deacetylation may indicate that both CDAs cooperate in chitin deacetylation of the outer cuticle layers. This is line with the results of a previous study, reporting that both TcCDAs are expressed in the same epidermal tissues, and yet depletion of either CDA alone resulted in cuticular defects. The authors suggested that the two proteins may cooperate with each other in chitin deacetylation [39]. For *TcCDA2* two splice variants (*TcCDA2a* and *TcCDA2b*) were reported with specialized functions in some tissues [39,42]. Since RNAi of either *TcCDA2a* or *TcCDA2b* affects cuticle integrity to various extents, we did not discriminate between both forms in our study and used dsRNA which silenced both splice variants.

In contrast, knockdown of midgut specific *TcCDA6-9* did not affect cuticle formation or chitin modification in the cuticle. The mKATE-CAP signal indicating the presence of low DA chitin was as strong as in the control animals.

Taken together, our observations suggest that CDA activity is required for the formation of the outer cuticular layer with low DA chitin and that both enzymes, TcCDA1 and TcCDA2, play a cooperative role in this process. The loss of Low DA chitin signals in response to RNAi-mediated *TcCDA1* and *TcCDA2* silencing is consistent with the findings of previous studies demonstrating that Tc*CDA1* and *TcCDA2* are essential for proper cuticle organization and chitin fiber assembly in insects [38,39].

## Conclusion

In this study, we combined X-ray diffraction, quantitative enzymatic mass spectrometry, and histochemical analyses to characterize the structural organization, abundance, and acetylation state of chitin in the cuticle and the peritrophic matrix (PM) of insects. XRD analysis of PM samples from different midgut regions of *Manduca sexta* revealed a high degree of structural similarity along the anterior–posterior axis. In all regions, the characteristic dihydrate β-chitin reflections were detected at comparable q-values, indicating that the chitin structure of the PM and its hydration state is uniform throughout the midgut [43,44].

The same diffraction characteristics were observed in whole-PM samples and in PM preparations from *Z. morio*, suggesting that β-chitin is probably conserved in the PM across lepidopteran and coleopteran species.

Quantitative enzymatic mass spectrometry revealed that chitin constitutes only a small fraction of the gut’s total mass, accounting for less than 1%. Moreover, gut chitin was found to be highly acetylated, with a DA of approximately 95% in control animals. Knockdown of gut-expressed chitin deacetylases resulted in a modest increase in acetylation, indicating that deacetylated chitin represents only a very minor fraction of gut chitin under normal conditions.

Consistent with these findings, histochemical analyses revealed strong staining for high DA chitin in the PM, whereas low DA chitin or chitosan could not be detected within the PM or in the gut epithelium under the conditions tested. In contrast, a clear stratification of chitinous material was observed in the larval cuticle, with low DA chitin present in the exocuticle and high DA chitin present in the endocuticle. After knockdown of either *TcCDA1* or *TcCDA2* low DA chitin disappeared in the exocuticle, demonstrating that both chitin deacetylases cooperate in chitin deacetylation. No comparable patterns of layered deposition of high and low DA chitin has been described in other insect species so far.

Overall, we identified β-chitin as the predominant allomorph present in the PM of lepidopteran and coleopteran insects and showed that PM’s chitin is highly acetylated. We further observed no differences regarding chitin allomorph and DA along the entire midgut. Future work will address how chitin–protein interactions, crosslinking, and higher-order matrix organization contribute to PM properties such as mechanical stability and permeability, and how these features are regulated during digestion and development.

## Acknowledgements

We thank Ahmet Sezer Bayrak for providing dsRNAs to silence *TcCDNA1* and *TcCDNA2*. We further acknowledge the Helmholtz-Zentrum Berlin (HZB) for the allocation of synchrotron radiation beamtime and hosting. X-ray diffraction data were collected at the mySpot beamline operated at the BESSY II synchrotron (Berlin-Adlershof, Germany). This work was supported by DFG grants ME2210/9-1 received in the scope of Joint Sino-German Research Projects 2021 and ME2210/11-1.

## Supplementary Information

**Figure S1.**
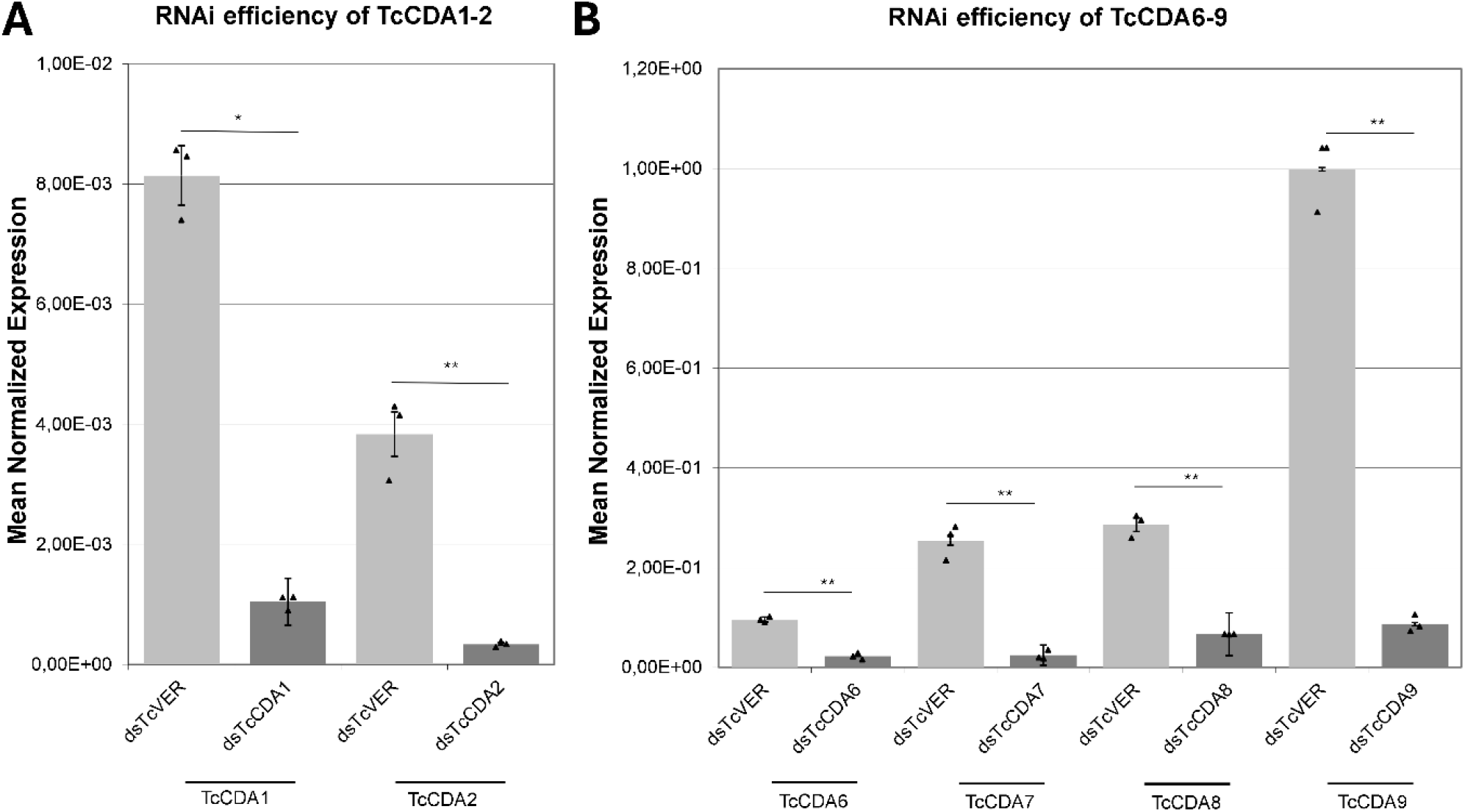
RNAi-mediated knockdown of midgut-expressed chitin deacetylases TcCDA6-9. **(A)** Relative expression levels of *TcCDA1* and *TcCDA2* in larval midguts following gene-specific RNAi to silence *TcCDA1* and *TcCDA2* compared to control larvae injected with *dsTcVER*. For the determination of RNAi efficiency, total RNA was isolated from one half of the larvae, while the tissue of the other half was used for histochemical analyses. **(B)** Relative expression levels of *TcCDA6, TcCDA7, TcCDA8* and *TcCDA9* following RNAi to silence their expression, compared to control larvae injected with *dsTcVER*. Expression levels were normalized to *TcRpS6*. Bars represent mean normalized expression (± SD) of biological triplicates (n=3), with individual data points shown. RNAi treatment resulted in strong reductions in transcript levels, corresponding to RNAi efficiencies of 87.2% (*TcCDA1*), 91.2% (*TcCDA2*), 76.8% (*TcCDA6*), 90.4% (*TcCDA7*), 76.8% (*TcCDA8*) and 91.3% (*TcCDA9*). Statistical significance was assessed using unpaired two-tailed Welch’s t-tests on ΔCt values. All knockdowns resulted in significant reductions in transcript levels either at *p* < 0.05 (*) or at *p* < 0.01(**).

**Figure S2.**
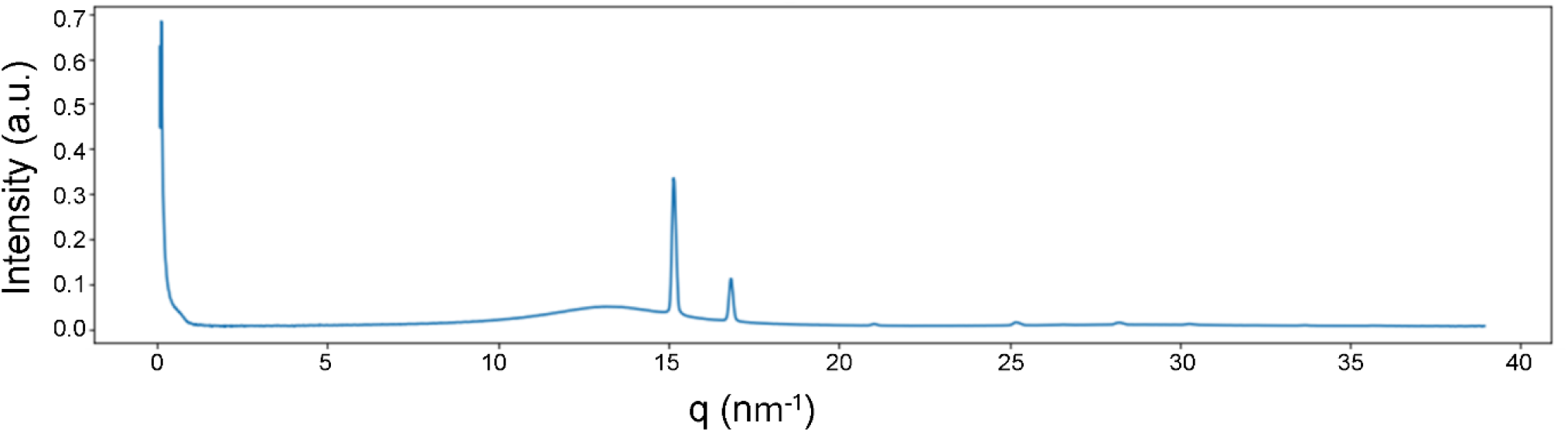
Representative XRD profile showing grease contamination. Example of a one-dimensional X-ray diffraction intensity profile displaying additional peaks originating from grease contamination. These features were present in a subset of measurements and are not related to the biological samples. While they did not affect the main analysis, the contaminated regions were excluded from further evaluation. This example is shown for transparency and documentation of data quality.

**Figure S3.**
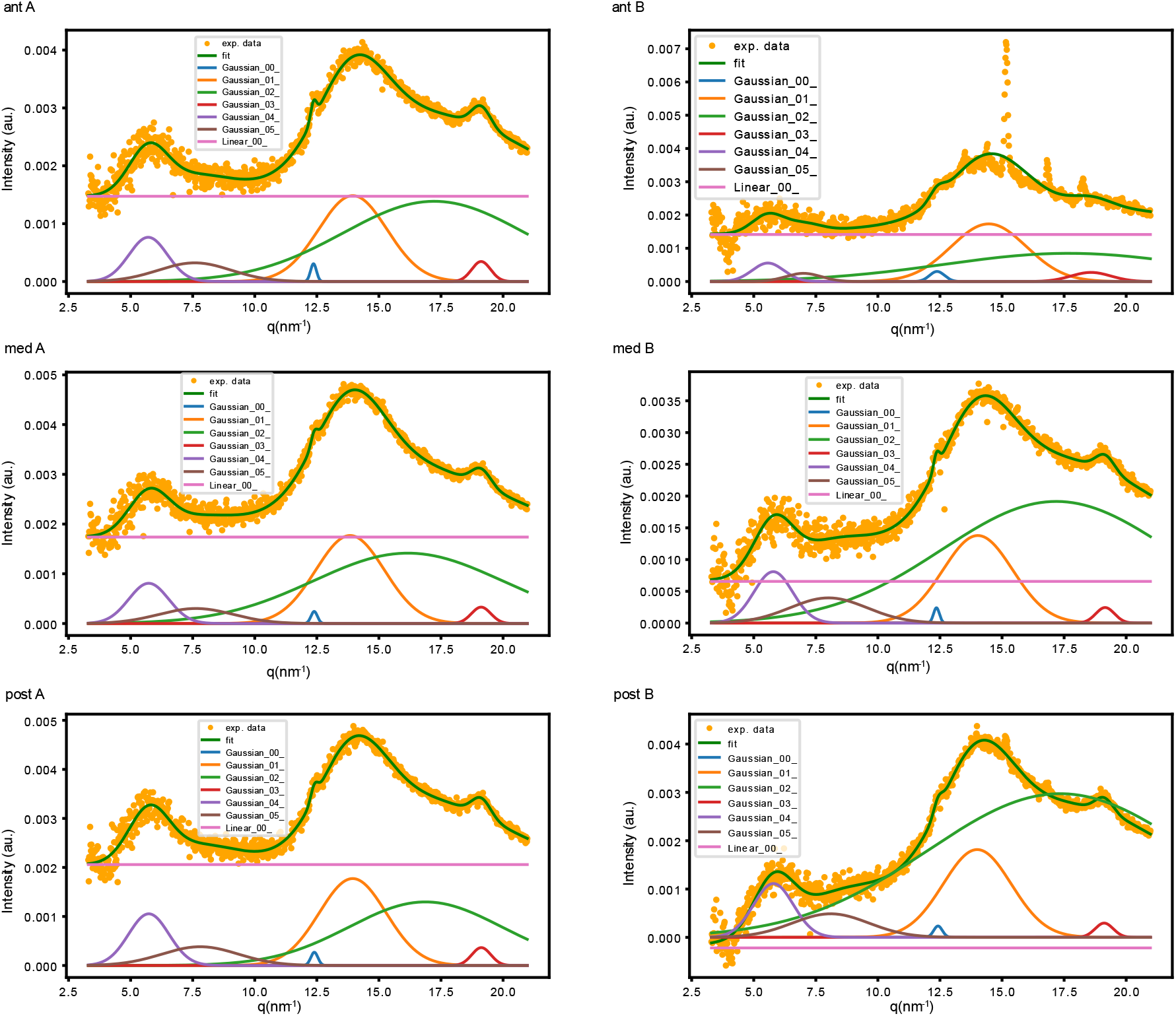
Gaussian peak fitting of X-ray diffraction profiles from regional PM samples of *M. sexta*. Azimuthally integrated one-dimensional diffraction profiles (orange dots) from anterior (ant), median (med), and posterior (post) midgut PM are shown for two biological replicates **(A**,**B)**. The experimental data were fitted using a multi-component Gaussian model (green line), consisting of contributions assigned to the (020), (110), and (013) reflections, as well as an amorphous background and linear baseline. Individual Gaussian components are shown as colored curves. The consistent peak positions and relative contributions across regions and replicates indicate a conserved structural organization of chitin within the PM.

**Figure S4.**
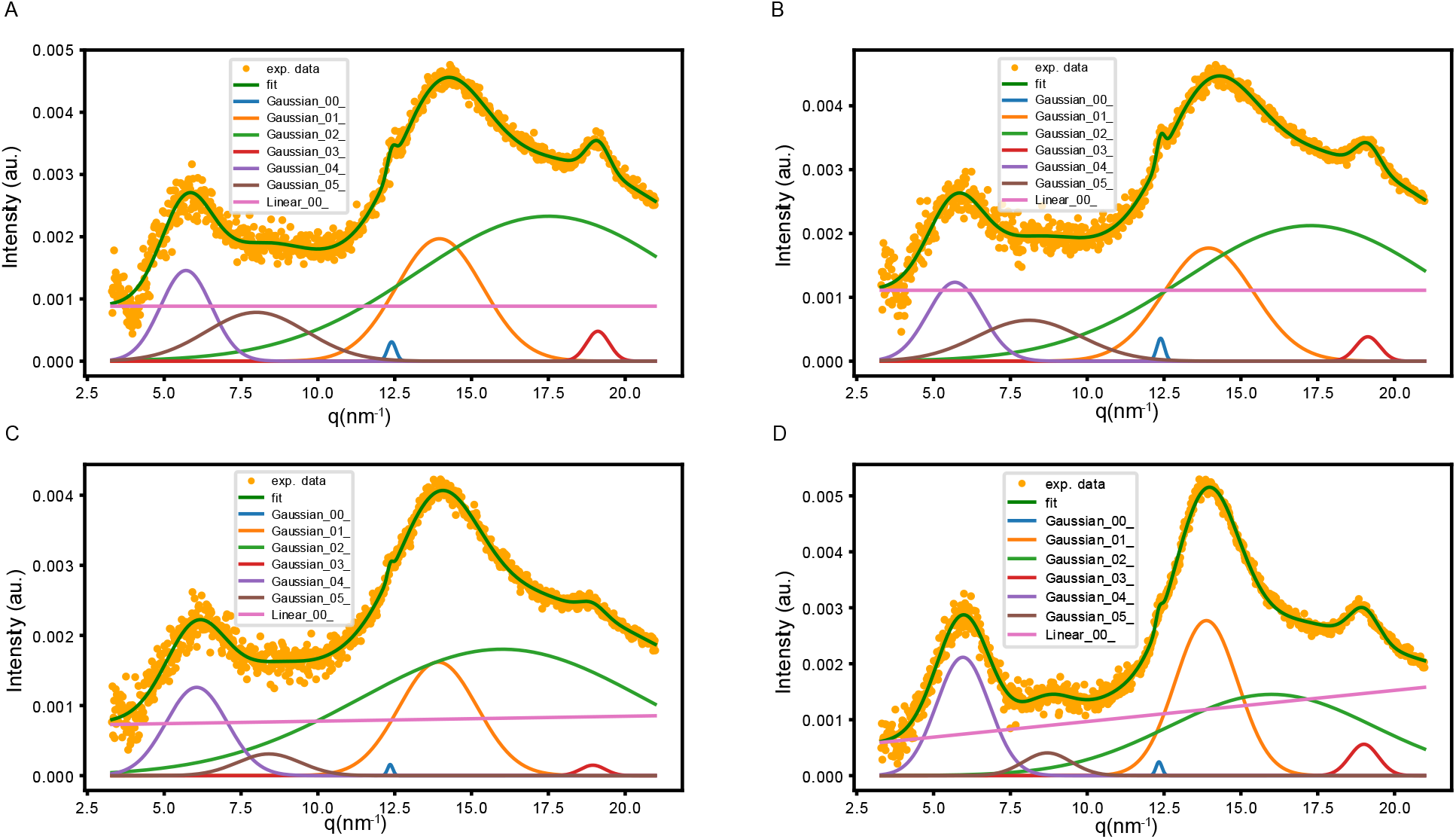
Gaussian peak fitting of diffraction profiles from whole PM samples of different insect species. Azimuthally integrated diffraction profiles (orange dots) of whole PM samples from *M. sexta* **(A**,**B)** and *Z. morio* **(C**,**D)** were fitted using a multi-component Gaussian model (green line). The fits include contributions corresponding to the (020), (110), and (013) reflections, as well as an amorphous component and baseline correction. Despite minor variations in peak shape and intensity, all samples exhibit a similar peak pattern, indicating a conserved β-chitin organization across species.

**Table S1.**
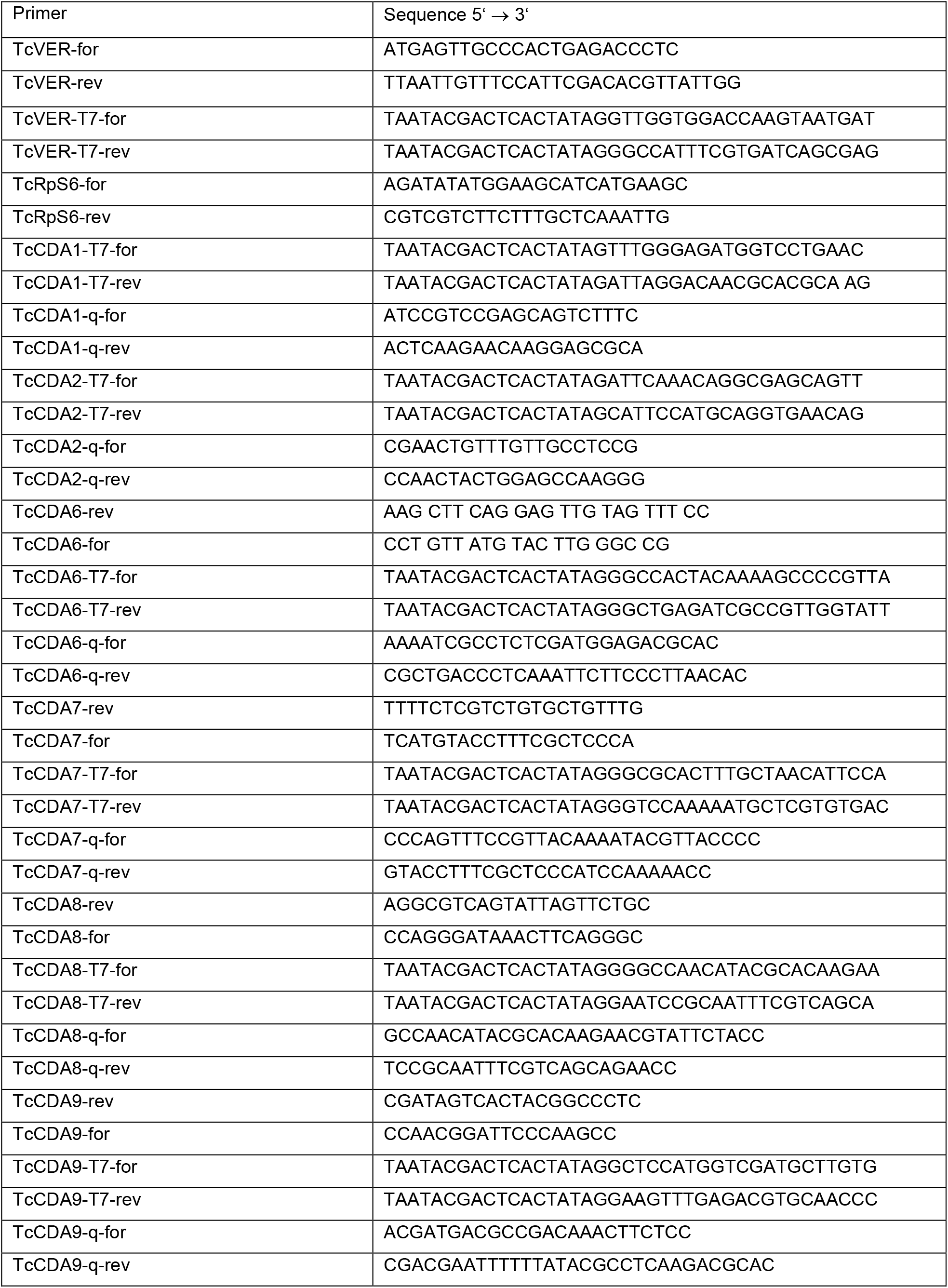
List of utilized primers.

**Table S2.** Gaussian decomposition parameters of X-ray diffraction profiles from different midgut regions of *Manduca sexta*. Parameters of multi-component Gaussian fits applied to azimuthally integrated diffraction profiles of anterior, median, and posterior PM samples. Listed are amplitudes, peak positions (center, q in nm^−1^), and peak widths (sigma), as well as linear background contributions. Median values were calculated across replicates to assess regional consistency of peak positions and relative contributions.

|  | Anterior A | Anterior B | Anterior (median) | Median B | Median C | Median (median) | Posterior B | Posterior C | Posterior (median) |
| --- | --- | --- | --- | --- | --- | --- | --- | --- | --- |
| Linear_00_slope | 0.000 | 0.000 |  | 0.000 | 0.000 |  | 0.000 | 0.000 |  |
| Linear_00_intercept | 0.001 | 0.001 |  | 0.002 | 0.001 |  | 0.002 | 0.000 |  |
| Gaussian_00_amplitude | 0.000 | 0.000 |  | 0.000 | 0.000 |  | 0.000 | 0.000 |  |
| Gaussian_00_center | 12.366 | 12.383 |  | 12.393 | 12.359 |  | 12.399 | 12.422 |  |
| Gaussian_00_sigma | 0.119 | 0.300 |  | 0.129 | 0.118 |  | 0.143 | 0.174 |  |
| Gaussian_01_amplitude | 0.005 | 0.007 |  | 0.006 | 0.005 |  | 0.006 | 0.006 |  |
| Gaussian_01_center | 13.948 | 14.465 | 14.207 | 13.839 | 14.025 | 13.932 | 13.944 | 14.000 | 13.972 |
| Gaussian_01_sigma | 1.379 | 1.515 |  | 1.384 | 1.406 |  | 1.346 | 1.374 |  |
| Gaussian_02_amplitude | 0.013 | 0.011 |  | 0.013 | 0.022 |  | 0.010 | 0.040 |  |
| Gaussian_02_center | 17.228 | 17.687 |  | 16.170 | 17.184 |  | 16.876 | 17.343 |  |
| Gaussian_02_sigma | 3.663 | 5.043 |  | 3.806 | 4.587 |  | 3.076 | 5.323 |  |
| Gaussian_03_amplitude | 0.000 | 0.001 |  | 0.000 | 0.000 |  | 0.000 | 0.000 |  |
| Gaussian_03_center | 19.117 | 18.582 | 18.849 | 19.117 | 19.152 | 19.135 | 19.125 | 19.113 | 19.119 |
| Gaussian_03_sigma | 0.380 | 0.800 |  | 0.390 | 0.375 |  | 0.382 | 0.364 |  |
| Gaussian_04_amplitude | 0.001 | 0.001 |  | 0.002 | 0.002 |  | 0.002 | 0.002 |  |
| Gaussian_04_center | 5.708 | 5.572 | 5.640 | 5.729 | 5.786 | 5.758 | 5.732 | 5.793 | 5.763 |
| Gaussian_04_sigma | 0.775 | 0.633 |  | 0.802 | 0.777 |  | 0.813 | 0.802 |  |
| Gaussian_05_amplitude | 0.001 | 0.000 |  | 0.001 | 0.002 |  | 0.001 | 0.002 |  |
| Gaussian_05_center | 7.584 | 7.000 |  | 7.635 | 8.012 |  | 7.801 | 8.097 |  |

**Table S3.** Gaussian decomposition parameters of X-ray diffraction profiles from whole PM samples of different insect species. Parameters of multi-component Gaussian fits for azimuthally integrated diffraction profiles of whole PM samples from *Manduca sexta* and *Zophobas morio*. The table includes amplitudes, peak positions (center, q in nm^−1^), and widths (sigma) for each Gaussian component, along with linear background parameters. Median values are provided to facilitate comparison of structural features across species.

|  | *M. sexta*<br>A | *M. sexta*<br>B | *M. sexta*<br>(median) | *Z. morio*<br>A | *Z. morio*<br>B | *Z. morio*<br>C | *Z. morio*<br>(median) |
| --- | --- | --- | --- | --- | --- | --- | --- |
| Linear_00_slope | 0.000 | 0.000 |  | 0.000 | 0.000 | 0.000 |  |
| Linear_00_intercept | 0.001 | 0.001 |  | 0.001 | 0.000 | 0.001 |  |
| Gaussian_00_amplitude | 0.000 | 0.000 |  | 0.000 | 0.000 | 0.000 |  |
| Gaussian_00_center | 12.400 | 12.388 |  | 12.353 | 12.335 | 12.900 |  |
| Gaussian_00_sigma | 0.138 | 0.125 |  | 0.087 | 0.098 | 0.300 |  |
| Gaussian_01_amplitude | 0.007 | 0.007 |  | 0.005 | 0.007 | 0.023 |  |
| Gaussian_01_center | 13.958 | 13.951 | 13.955 | 13.915 | 13.878 | 13.475 | 13.878 |
| Gaussian_01_sigma | 1.404 | 1.466 |  | 1.266 | 1.016 | 1.779 |  |
| Gaussian_02_amplitude | 0.025 | 0.022 |  | 0.021 | 0.012 | 0.016 |  |
| Gaussian_02_center | 17.511 | 17.279 |  | 16.000 | 16.000 | 17.585 |  |
| Gaussian_02_sigma | 4.317 | 4.116 |  | 4.656 | 3.332 | 3.788 |  |
| Gaussian_03_amplitude | 0.000 | 0.000 |  | 0.000 | 0.001 | 0.000 |  |
| Gaussian_03_center | 19.110 | 19.132 | 19.121 | 18.948 | 19.005 | 18.995 | 18.995 |
| Gaussian_03_sigma | 0.371 | 0.392 |  | 0.404 | 0.470 | 0.299 |  |
| Gaussian_04_amplitude | 0.003 | 0.003 |  | 0.003 | 0.005 | 0.003 |  |
| Gaussian_04_center | 5.709 | 5.696 | 5.702 | 6.056 | 5.954 | 5.792 | 5.954 |
| Gaussian_04_sigma | 0.828 | 0.886 |  | 0.991 | 0.855 | 0.995 |  |
| Gaussian_05_amplitude | 0.003 | 0.003 |  | 0.001 | 0.001 | 0.005 |  |
| Gaussian_05_center | 8.000 | 8.100 |  | 8.405 | 8.702 | 8.903 |  |

